# Characterization of carotenoid-producing *Muricauda* sp. strains isolated from the reef-building coral *Galaxea fascicularis* and subtropical seawater

**DOI:** 10.1101/2023.12.12.571358

**Authors:** Funa Endo, Keisuke Motone, Kenji Kai, Tomoya Kitakaze, Yumi Nishikawa, Yuna Nishimura, Ruriko Kitamura, Toshiyuki Takagi, Michihiro Ito, Natsuko Miura, Michihiko Kataoka

## Abstract

Bacterial symbionts in corals and coral-associated zooxanthellae are attracting increasing attention as potential probiotics. Some members of the family Flavobacteriaceae are zooxanthellae-associated bacteria, which are known to protect zooxanthellae from stresses such as heat and light by producing carotenoids that reduce reactive oxygen species production. This study characterized two Flavobacteriaceae bacteria, *Muricauda* sp. strains ORYM1 (NBRC115792) and ORYM2 (NBRC115793), isolated from the reef-building coral *Galaxea fascicularis* and its surrounding seawater in Okinawa, Japan, respectively. The *Muricauda* sp. strain ORYM2 was isolated from subtropical seawater, and carotenoid production was assessed using ORYM2 as well as ORYM1, which was previously isolated from *G. fascicularis. De novo* genome sequencing revealed that both strains contain complete sets of zeaxanthin biosynthesis genes, similar to those found in other *Muricauda* spp. Thin-layer chromatography and high-performance liquid chromatography analyses demonstrated that ORYM1 and ORYM2 produce several carotenoids. The bacterial strains and carotenoids identified in this study provide insights into the biological roles of zooxanthellae-associated bacteria in protecting zooxanthellae and reef-building corals from environmental stresses.

**Statements and declarations:** Competing interests: The authors declare no competing interests.

## Introduction

Corals play a critical role in maintaining biodiversity in subtropical marine ecosystems; however, they encounter severe threats due to global warming. To protect coral health, studies have increasingly focused on maintaining the well-being of corals under elevated seawater temperature and light stresses. Corals exhibit a unique symbiotic state with endosymbiotic microalgae (zooxanthellae of the family Symbiodiniaceae) and bacteria, collectively known as the coral holobiont (Bourne et al. 2016). Among these symbionts, bacteria play a crucial role in maintaining the healthy state of corals and zooxanthellae. Recent studies have shown that corals treated with a cocktail of coral-associated bacteria can act as coral probiotics, endowing corals with resistance against heat and light stresses and inducing metabolic and genetic changes (Peixoto et al. 2017, 2021; Santoro et al. 2021). Furthermore, a bacterial consortium has been developed to scavenge free radicals in cnidarians (Dungan et al. 2021). These bacterial cocktails typically contain specific bacteria that can effectively reduce light and heat stresses, such as bacteria that produce carotenoids, which are common antioxidants. For example, a marine bacterium of the family Flavobacteriaceae, GF1, which is most closely related to the genus *Muricauda*, has been shown to protect zooxanthellae from heat and high light stresses and may be useful as a coral probiotic (Motone et al. 2020). Moreover, the genus *Muricauda* is one of the core members of the bacterial family Symbiodiniaceae (Lawson et al. 2018).

Despite their potential usefulness, coral-associated carotenoid-producing bacteria have not been deposited in public repositories, and their general unavailability hinders progress in coral conservation research. Although carotenoid-producing bacteria have been isolated from seawater (Hameed et al. 2012; Yokoyama et al. 1994; Jiang et al. 2023), it remains unclear whether seawater-derived bacteria can be used as probiotics.

Previous studies have shown that bacteria similar to those found in corals also exist in the surrounding seawater (Kitamura et al. 2021). This study investigated whether *Muricauda* sp. strains could be isolated from subtropical seawater around corals in addition to corals and further characterized the carotenoid-producing capabilities of the isolated strains through *de novo* genome sequencing and carotenoid measurements. The obtained strains and information will contribute to investigating the role of *Muricauda* spp. in protecting zooxanthellae from environmental stress and incorporating these beneficial bacteria into the coral holobiont from seawater.

## Materials and methods

### Isolation of bacterial strains from seawater

Subtropical seawater was collected from Sesoko Island, Okinawa, Japan. The collected seawater was streaked onto marine agar medium (BD Difco Marine Agar 2216; Becton, Dickinson and Company, Sunnyvale, CA, USA), kept cool during transport to the laboratory in Osaka, Japan, and subsequently incubated at 25°C. The resulting colonies were inoculated into 5 mL of marine broth medium (BD Difco Marine Broth 2216; Becton, Dickinson and Company), which was shaken at 300 rpm and 25°C until bacterial growth was confirmed. These cultures were used for glycerol stock preparation and 16S ribosomal RNA (rRNA) sequencing.

### 16S rRNA sequencing

Bacterial genome extraction followed by 16S rRNA sequencing was conducted partially according to the previously described protocol (Kitamura et al. 2021). Bacterial cultures (400 μL) were centrifuged at 20,400 ×g for 5 min. The supernatant was discarded, and the cells were mixed with 100 μL of ddH_2_O and heated at 95°C for 5 min. After cooling on ice, the samples were centrifuged at 13,000 ×g for 1 min. The supernatants containing the genome were used to amplify the 16S rRNA gene using KOD FX Neo DNA polymerase (TOYOBO, Osaka, Japan) and the primers 27F (5′-AGAGTTTGATCCTGGCTCAG-3′) (Lane, 1991) and 1492R (5′-TACCTTGTTAGGACTT-3′) (Frank et al. 2008). Sanger sequencing of the amplified DNA fragments was performed by Eurofins Genomics, Inc. (Tokyo, Japan) using the same primers. The obtained DNA sequences were annotated using Nucleotide Blast (https://blast.ncbi.nlm.nih.gov/Blast.cgi).

### Phylogenetic analysis

Phylogenetic analysis was conducted using MEGA-X version 10.2.4 (Kumar et al. 2018) via the neighbor-joining algorithm and 1,000 bootstrap replications, based on 16S rRNA partial sequences.

### Extraction and purification of bacterial genomes

ORYM1 and ORYM2 genomes were extracted using Genomic DNA Buffer Set (Qiagen, Hilden, Germany) and Genomic Tip 20/G (Qiagen). The extracted genomes were assessed for concentration and purity using BioSpec-nano (Shimadzu, Kyoto, Japan) and Qubit™ 4 Fluorometer (Thermo Fisher Scientific, Waltham, MA, USA). To verify that no fragmentation occurred during extraction, the samples were also analyzed via 0.8% agarose gel electrophoresis (Nacalai Tesque, Inc., Kyoto, Japan).

### De novo *genome sequencing and analysis*

Library preparation and whole-genome sequencing of ORYM1 and ORYM2 were performed by Bioengineering Lab. Co., Ltd. (Sagamihara, Japan) using MGISP 960 (MGI Tech Co., Ltd., Kobe, Japan) and MGIEasy FS DNA Library Prep Set (MGI Tech) according to the manufacturer’s instructions. The reaction time for enzymatic cleavage was 4 min, and adapters from the MGIEasy DNA Adapters 96 (Plate) Kit (MGI Tech) were utilized. Circular DNA was prepared with the polymerase chain reaction product and MGIEasy Circularization Kit (MGI Tech) according to the manufacturer’s instructions. DNA nanoballs were created using the DNBSEQ-G400RS High Throughput Sequencing Set (MGI Tech), and sequencing was conducted on the DNBSEQ-400 (MGI Tech) platform with 2 × 200 bp. The obtained raw data were treated with Trimmomatic version 0.33 (Bolger et al. 2014) for adapter trimming and quality filtering. The fragment output was assembled using SPAdes version 3.15.3 (Bankevich et al. 2012).

### Annotation of zeaxanthin-producing genes

Assembled genomic sequences of GF1 (GenBank accession number PRJNA588666), ORYM1, ORYM2, and *Muricauda* sp. NBRC101325 (GenBank accession number SAMD00557617) were annotated using RAST (Aziz et al. 2008). Gene cluster comparison was performed using Clinker (Gilchrist and Chooi 2021).

### Carotenoid extraction from bacteria

Carotenoids were extracted from ORYM1 and ORYM2 using a modified version of the previously described method (Sakagami et al. 2010). Bacterial cells (∼60 mg, wet weight) were mixed with a 1:1 mixture of methanol and acetone (1 mL), and the mixture was vortexed for 30 min. After centrifugation at 15,300 × *g* for 5 min at 4°C, the supernatants were used for further analysis.

### Absorption spectral analysis

After transferring 150 μL of the crude carotenoid extract into a 96-well plate, absorption spectral analysis was performed using a microplate reader (SpectraMax 190; Molecular Devices, Sunnyvale, CA, USA).

### Thin-layer chromatography (TLC) analysis

For TLC analysis, a plastic TLC plate with a fixed layer of silica gel 60 (#1.05750.0001; Merck Millipore, Billerica, MA, USA) was employed, along with an expansion solvent containing hexane and acetone in a ratio of 7:4.

### High-performance liquid chromatography (HPLC) analysis

Carotenoid identification was performed using HPLC under the following conditions: COSMOSIL Packed Column 5C18-AR-II (250 × 4.6 mm, 5 μm; Nacalai Tesque); mobile phase, acetonitrile/methanol/chloroform (47:47:6); column temperature, 40°C; flow rate, 1.0 mL/min; detection wavelength, 450 nm; sample injection volume, 10 μL; and acquisition time, 15 min. Zeaxanthin dissolved in solvent (with the same composition as the mobile phase) was used as a standard at concentrations of 0.31, 0.47, and 0.94 μg/mL.

### Liquid chromatography–mass spectrometry (LC–MS) analysis

ORYM2 cultures were extracted with an equal volume of acetone for 1 day. The aqueous concentrate obtained by removing acetone under reduced pressure was applied to a Sep-Pak Plus C18 cartridge (Waters). After washing the column with 0.1% formic acid solution, the carotenoid components were eluted from the column using acetonitrile. The eluate was subjected to LC–MS analysis under the following conditions: InertSustain C18 column (150 × 2.1 mm, 3 μm; GL Sciences); eluent, 70%–100% acetonitrile in 0.1% aqueous formic acid; column temperature, 40°C; flow rate, 200 μL/min; sample injection volume, 5 μL, and detection electrospray ionization (ESI), positive.

### Availability of bacteria used in this study

*Muricauda* sp. strains ORYM1 and ORYM2 are available from the National Institute of Technology and Evaluation (Osaka, Japan), under accession numbers NBRC115792 and NBRC115793, respectively.

### Data availability

Raw sequencing datasets obtained via next-generation sequencing analysis of ORYM1 and ORYM2 genomes are available in the DDBJ Sequence Read Archive under the BioProject accession number PRJDB15463.

## Results

### *Isolation and draft genome analysis of* Muricauda *sp. strains ORYM1 and ORYM2*

*Muricauda* spp. isolated from the reef-building coral *Galaxea fascicularis* in a previous study (Kitamura et al. 2021) was named ORYM1. Table S1 presents the comprehensive list of bacteria isolated from seawater in this study. Among the 33 bacteria, only one bacterium was annotated as *Muricauda* sp., designated as ORYM2.

A summary of the draft genome analysis of ORYM1 and ORYM2 is illustrated in Figure 1a. The estimated genome sizes of ORYM1 and ORYM2 were 5.61 and 4.23 Mbp, respectively, comparable to the genome size of GF1 (5.28 Mbp) (Motone et al. 2020), a previously reported zeaxanthin-producing bacterium. Figure 1b shows the phylogenetic tree analysis of the 16S rRNA sequence of major *Muricauda* spp. and previously reported zeaxanthin-producing bacteria (asterisk). The zeaxanthin biosynthesis pathway was identified in the ORYM1, ORYM2, GF1, and NBRC101325 genomes (genome size: 3.88 Mbp). The *Muricauda* sp. strain NBRC101325, isolated from seawater in Yakushima Island (Kagoshima, Japan) by Kyowa Hakko Kogyo Co., Ltd., is available in live form from a public repository and is potentially useful as a carotenoid-producing bacterium. All four strains had zeaxanthin biosynthesis genes, namely, phytoene dehydrogenase, phytoene synthase, and β-carotene hydroxylase, clustered downstream of a MerR family transcriptional regulator (Fig. 1c). A lycopene β-cyclase gene, which catalyzes the reaction from lycopene to β-carotene, was also found in all four strains but not within the cluster (Table S2–S4). ORYM1, ORYM2, and GF1 genomes contained two phytoene dehydrogenase genes, whereas the NBRC101325 genome had only one gene, suggesting that ORYM1 and ORYM2 possess the ability for carotenoid biosynthesis, including zeaxanthin.

**Fig. 1.**
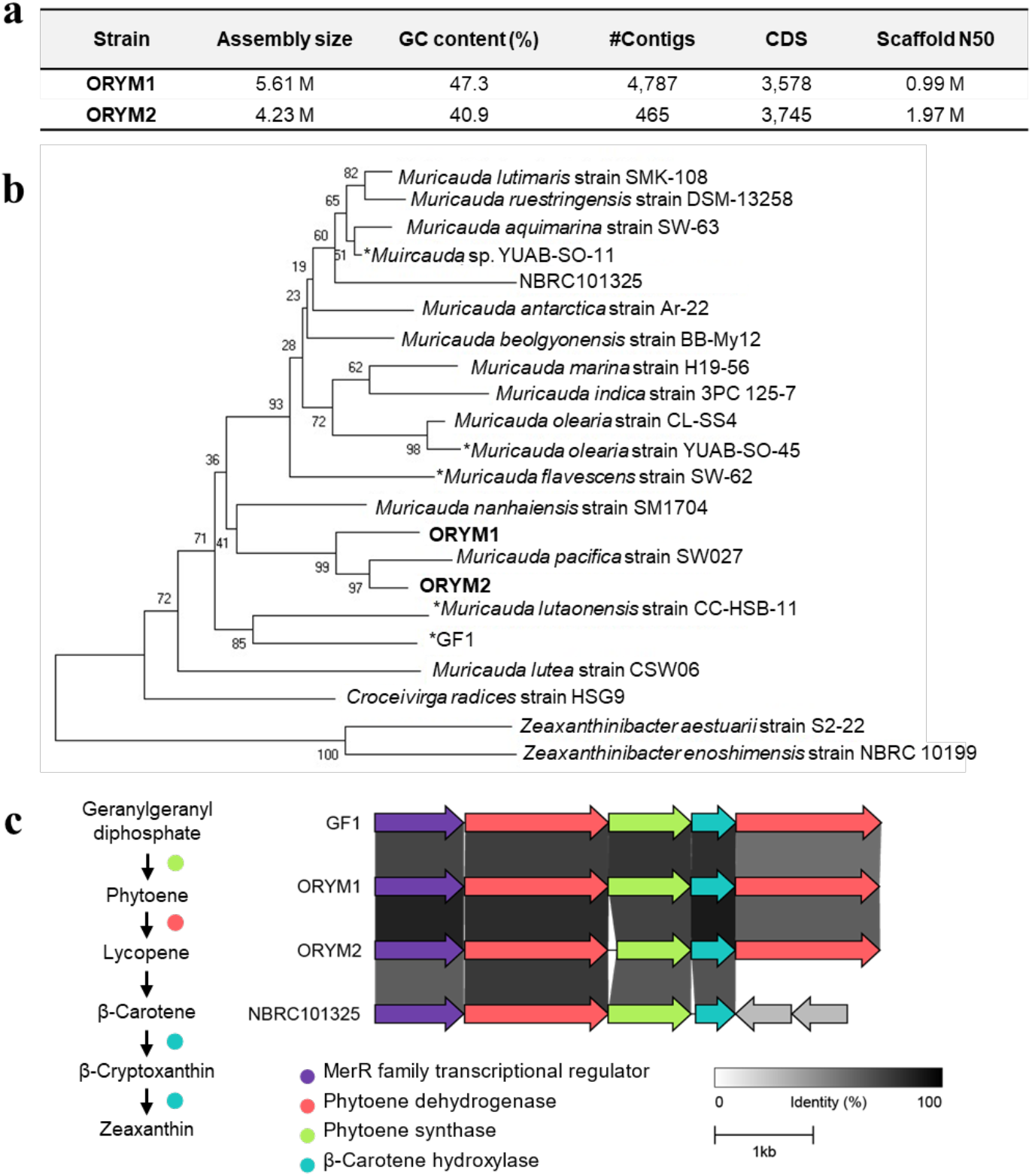
Characterization of *Muricauda* spp. used in this study. (a) Overview of the draft genomes of ORYM1 and ORYM2. (b) Phylogenetic analysis of 16S rRNA genes of ORYM1 and ORYM2. The asterisk indicates bacteria previously reported to produce zeaxanthin. (c) Comparison of carotenoid-producing gene clusters among *Muricauda* sp. strains GF1, ORYM1, ORYM2, and NBRC101325

### Analyses of carotenoid production by ORYM1 and ORYM2

To assess the carotenoid productivity in ORYM1 and ORYM2, their cell extracts were analyzed via ultraviolet-visible (UV-vis) spectrophotometry (Fig. 2a), TLC (Fig. 2b), and HPLC (Fig. 2c). The UV-vis spectra of the extracts exhibited absorption maxima at 470 and 500 nm (Fig. 2a), indicating the production of carotenoids by these strains. TLC and HPLC analyses indicated the presence of a compound that exhibits similar behavior as zeaxanthin and another major carotenoid with distinct chromatographic characteristics (Fig. 2b and c). To further characterize the major carotenoid, LC–MS analysis was performed (Fig. 2d). ESI–MS analysis of a major peak yielded ions at *m*/*z* 567 and 584 (Fig. 2d), which were inconsistent with the [M+H]^+^ value for zeaxanthin and other well-known carotenoids, indicating that the compound is an unknown carotenoid. Nuclear magnetic resonance analysis should be performed in the future to identify this compound in further detail.

**Fig. 2.**
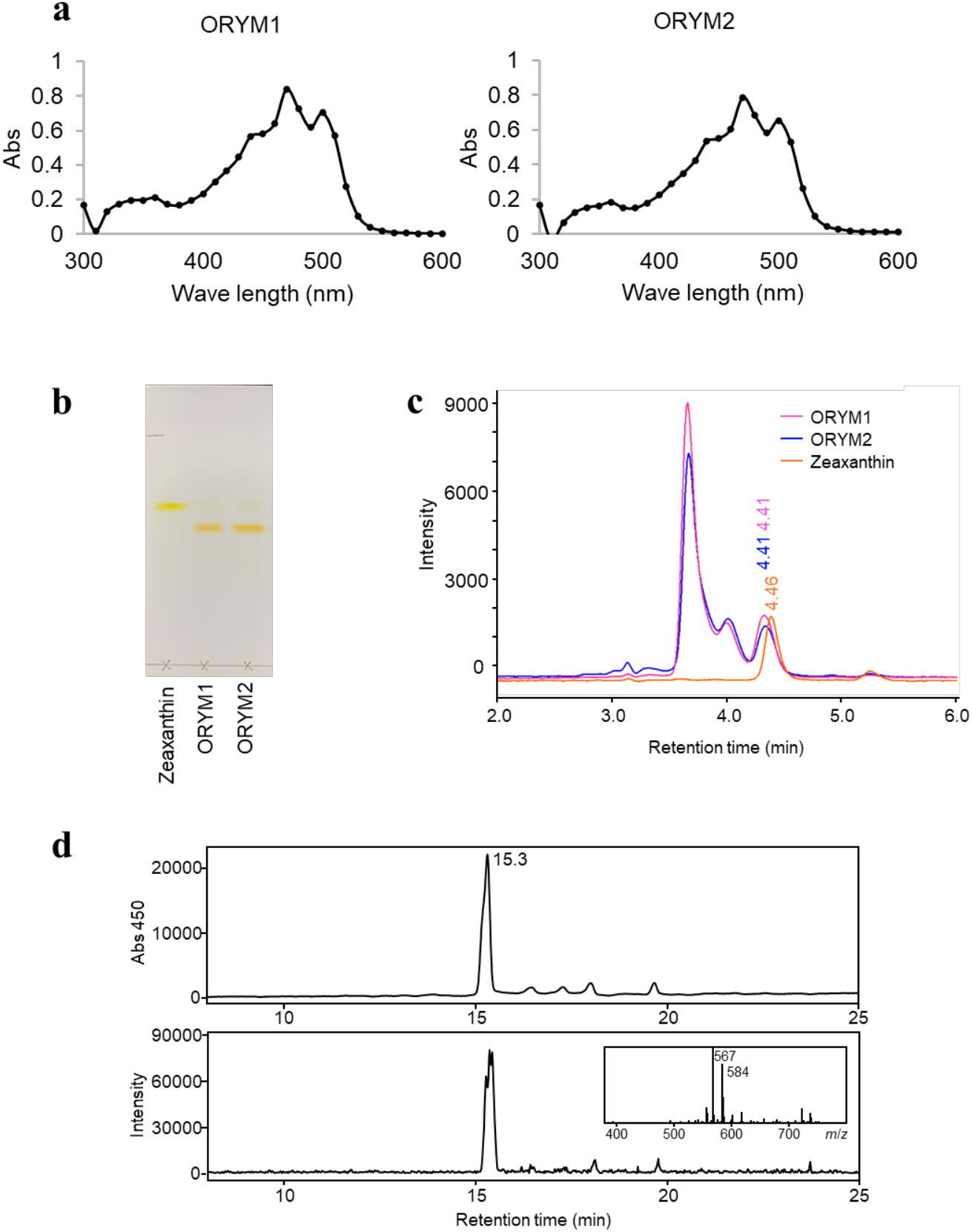
Carotenoid production by ORYM1 and ORYM2. (a) Absorption spectral analysis of crude carotenoid extracts (n = 3). Black dots indicate the average of three technical replicates. (b) TLC analysis. (c) HPLC analysis. Pink line: ORYM1; blue line: ORYM2; yellow line: zeaxanthin standard. (d) LC–MS analysis of ORYM2 cell extracts. Upper panel: a chromatogram of UV detection, lower panel: a chromatogram of SIM detection (*m*/*z* 567). ESI–MS data regarding a peak at 15.3 min are shown as an inset

## Discussion

This study successfully isolated carotenoid-producing *Muricauda* spp. from reef-building corals and subtropical seawater. Carotenoid-producing bacteria belonging to the genera *Muricauda* (Motone et al. 2020), *Maribacter*, and *Roseivirga* (Takagi et al. 2023) have attracted considerable interest as potential coral probiotics capable of protecting corals and their endosymbiotic microalgae from severe light and heat stresses. However, the diversity of carotenoids produced by coral-associated bacteria and their effectiveness in protecting microalgae, including zooxanthellae, remain poorly understood. This study will contribute to a better understanding of the role of bacteria in the reef ecosystem, particularly with regard to the acquisition of bacteria from the surrounding seawater and utilization of carotenoids by reef-building corals.

## Declarations

### Author contributions

Sample collection and isolation of bacteria: Funa Endo, Yumi Nishikawa, Yuna Nishimura, Ruriko Kitamura, Toshiyuki Takagi, Michihiro Ito, and Natsuko Miura; genome analysis: Funa Endo, Keisuke Motone, and Natsuko Miura; HPLC analysis: Funa Endo and Tomoya Kitakaze; LC–MS analysis: Kenji Kai and Funa Endo; writing—first draft: Funa Endo, Kenji Kai, Tomoya Kitakaze, Keisuke Motone, and Natsuko Miura; editing: Keisuke Motone, Kenji Kai, Tomoya Kitakaze, Toshiyuki Takagi, Michihiro Ito, Natsuko Miura, and Michihiko Kataoka; reviewing and approving the final manuscript: Funa Endo, Kenji Kai, Tomoya Kitakaze, Keisuke Motone, Yumi Nishikawa, Yuna Nishimura, Ruriko Kitamura, Toshiyuki Takagi, Michihiro Ito, Natsuko Miura, and Michihiko Kataoka.

### Ethics approval and consent to participate

Coral sampling in the previous study (Kitamura et al. 2021) was carried out with the approval of the authorities of Okinawa Prefecture, Japan.

### Data availability

Raw sequencing data sets obtained by next-generation sequencing analysis of ORYM1 and ORYM2 genomes are available in the DDBJ Sequence Read Archive under the BioProject accession number PRJDB15463.

